# Seq2DFunc: 2-dimensional convolutional neural network on graph representation of synthetic sequences from massive-throughput assay

**DOI:** 10.1101/2019.12.22.886085

**Authors:** Haotian Guo, Xiaohu Song, Ariel B. Lindner

**Affiliations:** INSERM U1284. Systems engineering and evolution dynamics, Paris, France; Center for research and Interdisciplinarity (CRI), Université de Paris, Paris, France

## Abstract

In recent years, a pipeline of massively parallel reporter assay (MPRA), and next-generation sequencing (NGS) provided large-scale datasets to investigate biological mechanisms in detail. However, bigger data often leads to larger complexity. As a result, theories derived from low-throughput experiments lose explanatory power, requiring new methods to create predictive models. Here we focus on modeling functions of nucleic acid sequences, as a study case of massive-throughput assays. We report a deep learning approach, training a two-dimensional convolutional neural network (CNN) on an ordered graph representation of nucleic acid sequences to predict their functions (Seq2DFunc). To compare the performance of Seq2DFunc with conventional methods, we obtained customized database on a CRISPR RNA processing assay. For this specific assay, analyses of sequence and RNA structure determinants failed to explain the results regardless of dataset size. 1-dimensional CNN of raw sequences generate generally failed to converge at < 10,000 or fewer sequences. By contrast, Seq2DFunc trained on ∼ 7,000 sequences still provided 86% accuracy. Given a sufficient dataset (∼ 120,000 sequences) for training, Seq2DFunc (96% accuracy, 0.93 f1-score) still outperformed the best 1D CNN (92% accuracy, 0.83 f1-score). We anticipate Seq2DFunc can be a versatile downstream tool for deciphering massive-throughput assays for many fundamental studies. In addition, the use of smaller dataset is especially beneficial to reduce the experiment budget or required sequencing depth.

## Introduction

Beyond encoding proteins as biological functions, nucleic acids play a critical role in direct regulation of fundamental biological processes. Non-coding sequences comprise a large fraction of human genomes (1), and other eukaryotic genomes (2). The majority of prokaryotic genomes also contains about 10% non-coding sequences (3). Even for the coding regions, the sequences are capable of regulating the expression and folding of proteins by modulating RNA structure and stability (4). However, the regulatory effects exerted by nucleic acid sequences also lead to difficulties in harnessing their potential in bioengineering. The causality between sequences and functions is mostly unclear, which makes the performance of designed genetic circuits hard to predict. Synonymous variants of GFP can result in 250-fold expression differences in *Escherichia coli* (5). Even the most basic elements such as promoters and ribosomal binding sites (RBS) will change their behavior in response to the sequence of other parts in the synthetic genetic circuits (6).

The regulatory mechanisms of nucleic acids usually present a high-degree of complexity. As a consequence, low-throughput experiments and high-throughput screening of 100 - 1,000 samples are often insufficient to unveil the design principles underlying the genotype to phenotype relationships. For instance, models of ribosomal binding site (7, 8), terminator (9), riboswitch calculator (10), though successfully predict their strengths, possess merely a relatively low explanatory power around 40 - 70% (7 - 10). In recent years, the method of massively parallel reporter assays (MPRA) was established to characterize sequence variants on an unprecedented scale from 6,500 up to 100 million sequences (14 - 24). FACS-seq is a commonly-used MPRA (14, 16, 18, 21, 24), in which elements of interest regulate the expression of fluorescent protein so that the measurement of entire library can be achieved by fluorescence-activated cell sorting (FACS) followed by next-generation sequencing (NGS). Other MPRA methods measure the elements by cell growth selection (19, 21), RNA copy number (15, 17, 20, 23), ribosome profiling (22) etc. for easy incorporation with downstream NGS. Especially, thanks to nowadays advances in DNA synthesis technology (25), it is possible to measure not only random sequences, but also rationally libraries designed for specific queries and hypotheses (14, 15, 17, 19 - 22). The combination of MPRA and NGS have been used in characterizing the synthetic 5’ untranslated region (5’-UTR) (16, 19, 21, 22), 3’-UTR (20), promoters (14, 24), ribozyme (18, 23), and human cis-regulatory DNA elements (15, 17).

The larger dataset sometimes can result in new hurdles in the analysis. The biological mechanisms underlying nucleic acid sequences are seldom governed by one or a few dominant laws, but rather by a list of entangled features (21). It seems unlikely to be solved by simple models. Therefore, powerful tools are required to decipher the logic behind all the four-letter codes. Many analyses we use today were developed in the era of low-throughput experiments, and may fail to explain the intricacy in the data from massive-throughput assay such as MPRA-NGS pipeline. In recent years, researchers applied the technique of deep learning, such as convolutional neural networks (CNN) on sequencing data (Figure 1A) (19, 22, 26, 27). These CNNs are good at extracting local patterns and motifs in 1-dimensional, linear sequence, and are capable to learn useful global interactions with higher-order convolutions (28). Nonetheless, the function of nucleic acid usually requires long-distance interactions (29, 30), which are not a global feature nor sequence motif, but local pattern in 2-dimensional space. Hence, we propose a deep learning model, to map nucleic acid **Seq**uences to **Func**tions by a **2-D**imensional CNN on the graph representation of sequences, Seq2DFunc. In the Seq2DFunc, all possible interactions between bases are identified, to create matrices that allow a 2-dimensional CNN to identify 2-dimensional neighbours, without losing 1-dimensional information.

**Figure 1.**
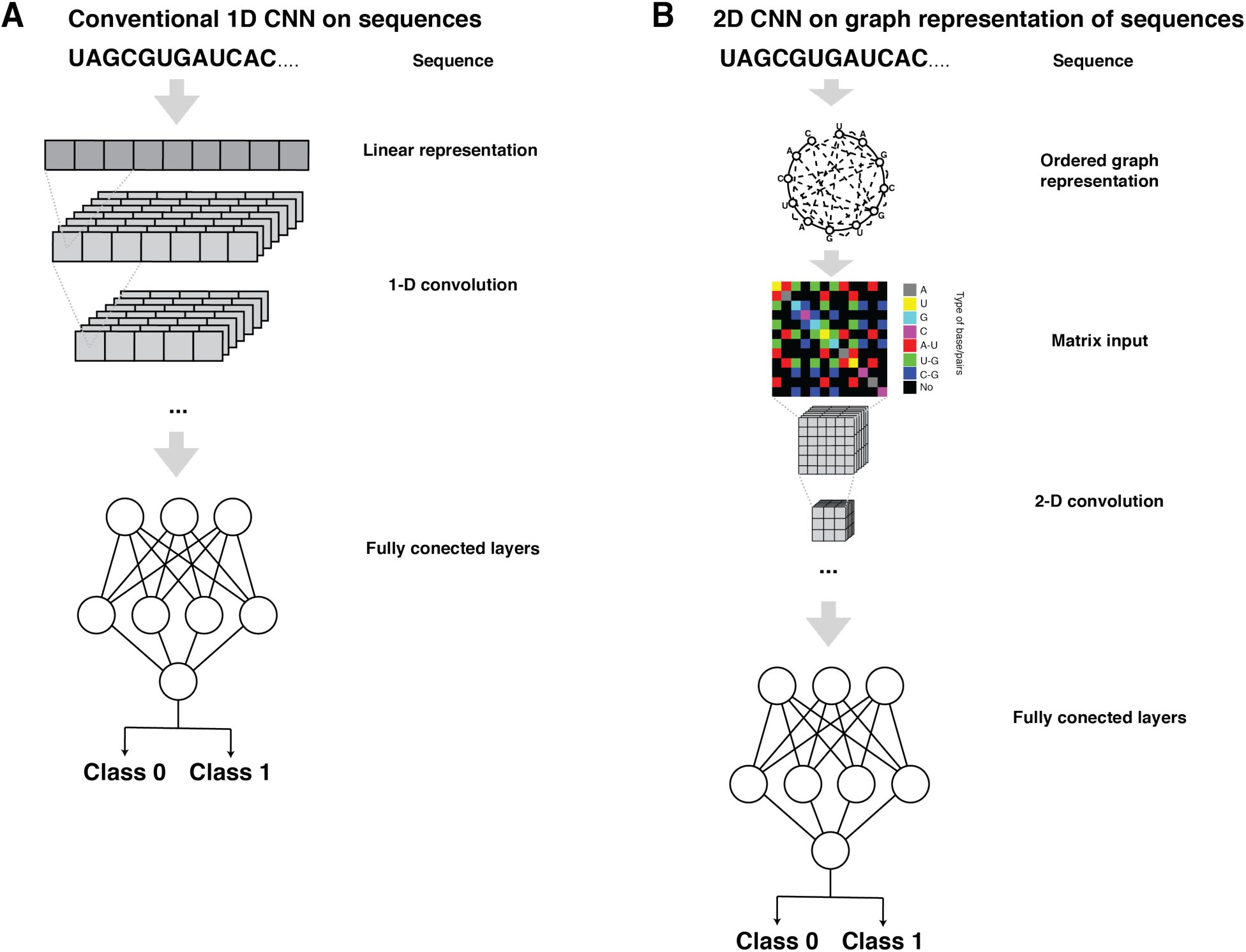
Architectures of 1-dimensional convolution neural network (A), and Seq2DFunc (B). In 1D CNN (A), sequences are represented by a linear vector, and then processed by 1-dimensional convolution. By contrast, in Seq2DFunc (B), sequences are represented by an ordered graph first, and then a 2-dimensional matrix. 2D matrices are processed by 2-dimensional convolution. In both 1D CNN and 2D CNN Seq2DFunc, fully connected layers take the outputs from convolutional layers, to generate the final prediction, as class 0 or 1 for example.

To validate the rationale, we used an in-house unpublished database on CRISPR RNA processing. Traditional methods including k-mer search, thermodynamic analysis, and kinetic folding failed to explain the observation, individually, or together in a machine-learning model. 1D CNN failed to converge, when it’s trained with < 10,000 sequences. In contrast, Seq2DFunc could be trained on a smaller dataset (∼ 7,000) and perform with 86% accuracy. When dataset size is increased, Seq2DFunc provided over 96.04% accuracy and 0.9319 f1-score, again outperforming 1D CNN results with 91.83% accuracy and 0.8308 f1-score. We hope this Seq2DFunc can be a useful alternative to the current methods for the analysis of the enormous datasets produced by massive-throughput assays, to build highly accurate predictive models. Furthermore, the requirement of smaller dataset is specific favorable as the cost of data point is still expensive.

## Results

### Design of Seq2DFunc

First, we span 1-dimensional sequence information into 2 dimensions using graphs (Figure 1B). For a N-nucleotide sequence, we consider an ordered graph with N nodes. Each node represents the respective base in the sequence, and is labeled by its content, as A, T (or U), G, and C. Then, all the possible interactions between two nodes are added as edges of the graph. Here, rather than simply applying a complete graph we applied prior knowledge to help the training process. In the following study of RNA elements, we assumed the main pairwise interaction comes from the RNA structure, and thus allowed only Waston-Crick base pairings, A-U, G-C, and a non-canonical base pairing G-U (31). This choice was guided by the trade off between the efficiency of training, and the potential loss of the information, and depends on researchers’ objective - for instance, higher-order structure such as G-quadruplex will not be represented if only 3 types of base pairs are allowed. Following this, we created a N-by-N matrix, by adding adjacency matrix indicating all the edges, and a diagonal matrix for all the nodes. To be noticed, the matrix is symmetric.

For further computation in deep learning, N-by-N matrix is then encoded by a method of choice. A commonly used approach is one-hot encoding (32) (see Method). However, the resulted N-by-N-by-8 array can be too sparse for some algorithms. Alternatively, we also attempted label encoding (or integer encoding), turning base content and base-pairing types into integers, in a specifically chosen arrangement (see Method). The encoded arrays are used as the input of a 2-dimensional CNN that uses 2-dimensional convolutional kernels to capture the local patterns. To handle imbalanced dataset from most of biological measurements, it’s also possible to choose data to create a smaller set that are uniformly distributed for training, but such dataset size may not be large enough to train a neural network, and will lead to biases, and information loss. To overcome this issue, we use the cross-entropy loss function for better training (33).

### Example database on CRISPR RNA processing without obvious classic features

To compare Seq2DFunc with other methods that have been used in previous studies on MPRAs, we used an in-house database on CRISPR RNA processing from MPRA (unpublished). Large libraries of random sequences were synthesized and characterized by FACS-seq to decipher how they influence the efficiency of CRISPR RNA processing mediated by *Pseudomonas aeruginosa* Cas6f, Csy4. Each sequence was classified as 0 or 1 according to the activity (see Method). We chose this dataset as our challenge because it represents a class of non-trivial RNA structural dependencies (21) that conventional analysis mostly failed to explain.

We first applied k-mer searches to investigate if any sequence motifs occur. k-mer abundances (k from 2 to 4) are compared between the two categories. In each analysis, the frequencies of each k-mer in two categories are highly correlated, with correlation coefficients as 0.78, 0.70, and 0.62 respectively (Figure 2A). None of the k-mers could be used as a predictive indicator (Figure 2B).

**Figure 2.**
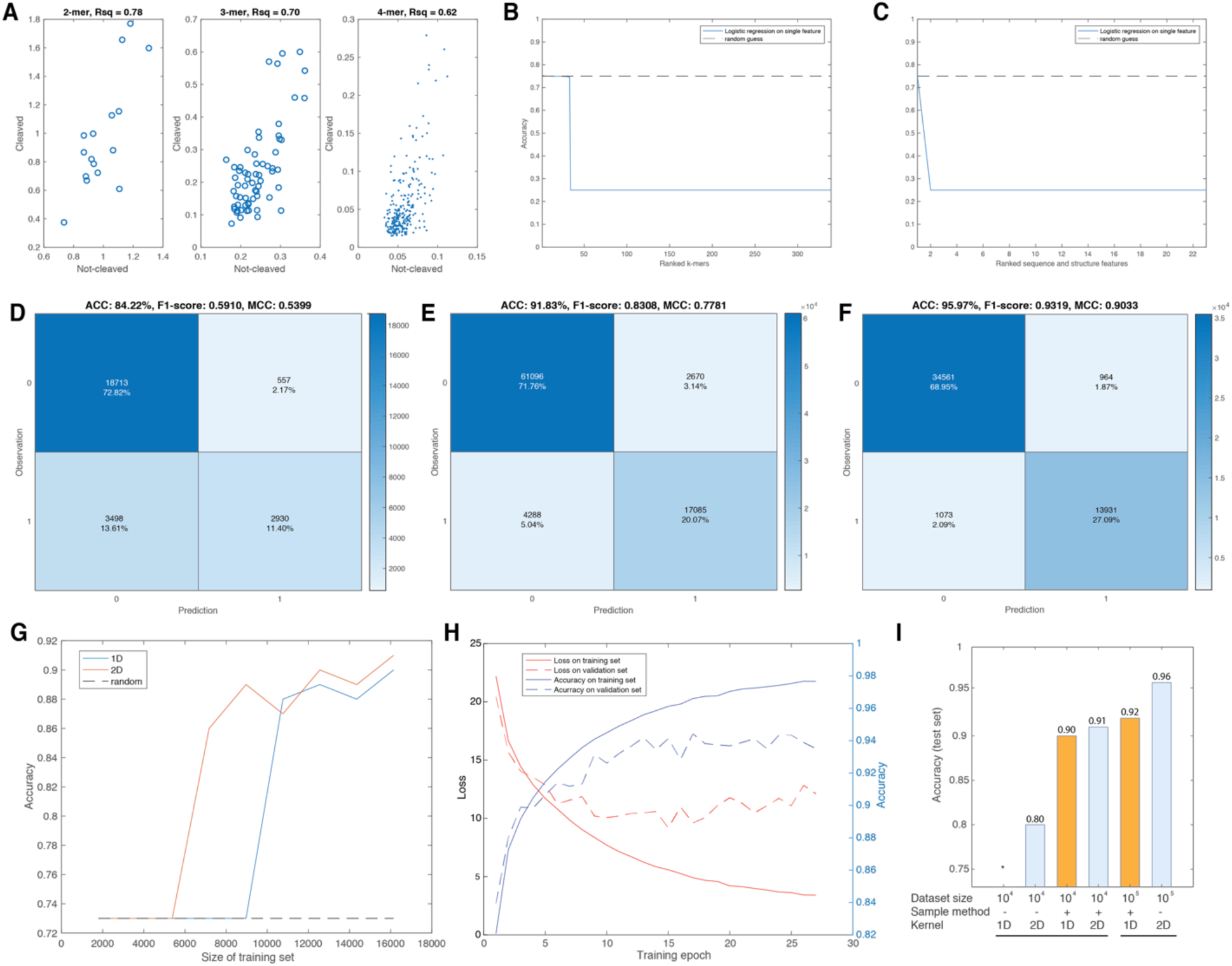
Seq2DFunc outperforms the conventional analysis and 1D CNN. A. The probability of k-mers (k as 2, 3, 4) detected in not-cleaved (class 0), and cleaved (class 1) are highly correlated while no significant outlier is identified, implying the frequency differences of k-mers between two classes are random errors. B. None of k-mers can be used to predict the class of sequences. Accuracy of the logistic regression models trained on individual k-mers were ranked and shown in blue curve. As a comparison, random guess was shown as a black dashed line. Prediction using k-mer is not significantly better than a random guess. C. Other analysis including thermodynamics, kinetics, and codon biases features, provided similarly poor predictive power. D - F, Confusion matrices of the best random forest model trained on the features shown in C, the best 1D CNN model, the best Seq2DFunc model. The size of training sets are all set as 70% of the large dataset containing >100,000 sequences. Confusion matrices showed the prediction versus observation on the test sets that were unseen in the training. The size of test sets varied due to the sampling methods (see Methods). Accuracy (ACC), f1-score, and mathew’s correlation coefficient (MCC) were calculated for each confusion matrix. G. Accuracy on unseen test set of 1D CNN and Seq2DFunc models trained on training sets across various sizes. Random guess was shown as a black dashed line as a baseline. H. Convergence of Seq2DFunc trained on small dataset (∼10,000) without bootstrap sampling method. I. Summary of neural network models in different training conditions. 1D CNN are labeled in yellow, and 2D CNN Seq2DFunc are labeled in blue. Models trained on small dataset are labeled as 10^4, and the large dataset as 10^5. Bootstrap sampling are marked as “+”, and simple random sampling as “-”. In all models trained on small dataset, 1D CNN with random sampling have no training succeeded (marked as *), 2D CNN Seq2DFunc with random sampling reached 80% accuracy, 1D CNN with bootstrap sampling reached 90% accuracy, and 2D CNN Seq2DFunc with bootstrap sampling reached 91% accuracy. In all models trained on large dataset, the best 1D CNN was trained with bootstrap sampling and reached 92% accuracy, and the best 2D CNN Seq2DFunc was trained with simple random sampling and reached 96% accuracy.

We then used NUPACK (11) to calculate thermodynamics of RNA secondary structures, including partition function, complex energy, minimum free energy (mfe), mfe structure, probability of mfe structure, and the total number of possible structures (see Method). The mfe structure was used for detailed analysis. We checked whether Csy4 binding site folded into the correct stem-loop structure. Then we segmented mfe structure into smaller fragments, and count the number of bases included (see Method). Using Kinwalker (13) from ViennaRNA (12) packages, we predicted the RNA folding trajectory (see Method). The intermediate structures, energy barrier and time required to get into the next structure, and running time were recorded for each sequence. The statistics including mean, median, standard deviation, and maximum were applied on the energy barriers of each trajectory. Number of total intermediate structures were also counted. Because the reporter assay involved protein translation, we also evaluated the codon usages, including codon fraction in the same amino acid, and the codon frequency in 1,000 codons of all amino acids (34).

Unfortunately, none of the features from above analysis was capable to classify two categories either (Figure 2C). Next, we sought if as a whole, all the features can be used to predict the classes. We trained a random forest (35) using all the computed features as input (see Method). Evaluated on the unseen test data, the random forest model was capable to identify the more abundant category, but was almost randomly guessing when predicting the other one (Figure 2D).

### 1D CNN is capable to extract the features but only with big enough data

Given that the sequence and structural analysis were largely inefficient for this specific dataset, we used 1D CNN as the baseline method. We first applied 1D CNN on a library of ∼120,000 sequences. Using bootstrap sampling to improve the training set, we obtained the best 1D CNN model reaching 91.83% accuracy on test sets (Figure 2E, see Method). 1D CNN could be trained without bootstrapping, but performed worse (data not shown). Then, we asked whether 1D CNN can work on a smaller dataset. To this end, we trained 1D CNN model from scratch on a smaller dataset from an independent experiment. We found 1D CNN cannot converge on this dataset without bootstrapping (Figure 2I). Using bootstrap sampling method to augment the data, the accuracy was higher than a random guess only when >10,000 sequences were used for training (Figure 2G). Thus, 1D CNN is capable to extract the features underlying the mechanism MPRA characterized, but only when the dataset is sufficient.

### RNA Seq2DFunc outperforms 1D CNN on large library and requires fewer data for training

After accomplishing the baseline deep learning model, we trained Seq2DFunc on our RNA sequence data, i.e. RNA-Seq2DFunc. We obtained a better model with 96.04% accuracy (Figure 2F), outperforming 1D CNN. Also, 2D CNN RNA-Seq2DFunc showed a more balanced precision and recall, with f1-score 0.9319, significantly higher than 1D CNN with f1-score 0.8303. In addition, bootstrapping is not used in this model, and found redundant when sufficient data are available, although it is required to reach high accuracy for 1D CNN model.

We then tested 2D CNN RNA-Seq2DFunc on smaller training sets, as we did before on 1D CNN, to estimate the efficiency of RNA-Seq2DFunc on extracting the features from data. Though 1D CNN failed to work without bootstrapping technique, RNA-Seq2DFunc can converge and performed > 90% accuracy on the validation set (Figure 2H). When bootstrapping was applied, 2D CNN RNA-Seq2DFunc trained on only 7,171 sequences can provide over 86% accuracy (Figure 2G), which is still ⅔ of minimal requirement for training 1D CNN. Thus 2D CNN RNA-Seq2DFunc can outperform 1D CNN when the mechanistic rationale is intricate as in this example, and it may require much fewer data and simpler methods to be trained into an accurate enough model.

## Discussion

Synthetic biology provides a great opportunity to study the fundamental rules of biology. By blending experimental, data-driven approach with theoretical analysis, we learn what is true by what is made. The combination of massively parallel reporter assays and NGS leads to large datasets with a relatively low cost per measurement. It enables in principle to unveil phenomena that are intangible under low-throughput experiments, yet calls for new methodologies to understand the massive data produced. In this study, we demonstrated the shortcomings of traditional methods, and the more recent 1-dimensional CNN approach. Based on our prior understanding on the nature of nucleic acids’ regulatory function, we present a new 2D representation method of nucleic acid sequences and coupled it with a2-dimensional CNN, Seq2DFunc. On an unusually complex dataset of CRISPR RNA processing, we compared 2D CNN RNA-Seq2DFunc to the conventional analysis of RNA sequences and structures, machine learning on features, and 1-dimensional CNN on sequences. 2D CNN RNA-Seq2DFunc model outperforms all these methods, and requires a significantly smaller training dataset to converge as compared to 1D CNN. Importantly, the graph representation does not add new information, but only represents the sequence data in a higher dimension. We speculated that Seq2DFunc is more efficient because pairwise local patterns become easier to extract. Although only RNA dataset was tested in this study, we anticipate the same method can be applied for non-coding DNA sequences as well. 2D CNN Seq2DFunc also includes more parameters than 1D CNN by default, so we cannot ruled out that the training process benefited from the changing structure of neural networks.

We found that 2D CNN Seq2DFunc can be trained on 10-fold fewer data than 1D CNN. Even with techniques such as bootstrapping to augment the data, 1D CNN still requires 50% data in training to perform at the same accuracy. This is specifically advantageous for research in biology because the total cost to acquire enough data is still more expensive than that in information technology. According to our experience, an MPRA-NGS experiment containing one control, and one treatment group, with 3 replicates each, costs ∼ 0.05 euro per valid data points. For 1D CNN and other traditional analyses, 0.5 million to even 100 million sequences were required (19, 22, 24). In this study, when only 33-nucleotides are involved, the simple method of 1D CNN still needed over 100,000 sequences for training. Thus we expect that the cost for a thorough analysis, is from 5,000 - 25,000 euro at least. If a library of 0.5 million rationally designed sequences is characterized, the cost can even go up to ∼ 70,000 euro. The efficiency of Seq2DFunc allows the possibility to use fewer data, which can dramatically reduce the cost per experiment.

As the drawbacks of Seq2DFunc, it first requires a transformation of the sequence data. The time complexity increases quadratically in response to the length of sequences. According to the “no free lunch theorem” (36), we do not expect Seq2DFunc will be a final solution for all scenarios of nucleic acid sequence analysis. In some cases, it is very possible that local structural motifs, 1-dimensional sequence motifs or other features are dominant and easy to be extracted by traditional methods, 1D CNN or recurrent neural network (RNN). In such cases, applying Seq2DFunc may not provide a higher accuracy, or marginal effect is negligible. Further analysis of the neural network, and testing on other datasets are still needed for better understanding the pros and cons of this method.

In summary, we anticipate Seq2DFunc can be a powerful alternative to the existing methods on theoretical analysis of big data produced by MPRA, and provide further insights in complex regulations mediated by nucleic acids. We also hope to encourage the scientific community to adopt this MPRA-NGS-Seq2DFunc pipeline on the topic of interest for the future optimization of the deep learning approach in biology.

## Supporting information

Materials and Methods, Data and code availability

## Acknowledgement

We thank Yann Ponty’s advice on RNA structure analysis, David Bikard’s advice on CRISPR biology, and their inputs on deep learning section. We also thank colleagues at INSERM U1001 and CRI, and members in Ponty group from Ecole Polytechnique for the discussion on the project. This project has received funding from the European Union’s Horizon 2020 Research and Innovation Programme under the Marie Sklodowska-Curie Grant Agreement No. 665850.

## Attribution

H.G conceived the Seq2DFunc, provided the database on CRISPR RNA processing, and performed conventional analysis and random forest models, and wrote the manuscript. X.S performed the 1D CNN and Seq2DFunc, and provided the related methods for the manuscript. H.G and X.S analyzed the outcomes of analyses and models. A.B.L supervised the study and edited the manuscript.

