## Supplementary material for "Seq2DFunc: 2-dimensional convolutional neural network on graph representation of synthetic sequences from massive-throughput assay": Materials and Methods, Data and code availability

This PDF file includes:

Materials and Methods

Data and code availability

### Materials and methods

#### Deep learning architectures

For the one-hot encoding, bases A, U, G, C are encoded as [1,0,0,0], [0,1,0,0], [0,0,1,0], and [0,0,0,1]. For the label encoding, bases A, U, G, C are encoded as 1, 2, 3, 4, and base pairings A-U or U-A, U-G or G-U, G-C or C-G, as 5, 6, 7. This specific assignment are arbitrarily assigned due to the fact that U and G are capable to pair two types of bases.

The size of training set is 70% of the total data, unless it's specifically tuned to find the response between accuracy versus training set size. Two methods were applied: (1) Bootstrap sampling: Sequences in the training set is randomly sampled with replacement. Sequences not chosen (~50% of database) are used as test set. (2) Simple random sampling: Sequence in the training set is randomly sampled without replacement. Thus all sequences in the training set is unique. Sequences not chosen (30% of database) are used as test set.

Both methods were tested. The best model of 1D on the large dataset is trained using bootstrap sampling. The best model of 2D model on the large dataset is trained using simple random sampling.

For the 1D CNN, the architecture of network is as the following:

| Layer (type) | Output Shape | Param # |
|---|---|---|
| embedding_1 (Embedding) | (None, 31, 8) | 512 |
| conv1d_1 (Conv1D) | (None, 29, 128) | 3200 |

| | | |
|---|---|---|
| conv1d_2 (Conv1D) | (None, 27, 128) | 49280 |
| max_pooling1d_1 (MaxPooling1 | (None, 13, 128) | 0 |
| conv1d_3 (Conv1D) | (None, 11, 256) | 98560 |
| conv1d_4 (Conv1D) | (None, 9, 256) | 196864 |
| max_pooling1d_2 (MaxPooling1 | (None, 4, 256) | 0 |
| conv1d_5 (Conv1D) | (None, 2, 512) | 393728 |
| global_average_pooling1d_1 ( | (None, 512) | 0 |
| dropout_1 (Dropout) | (None, 512) | 0 |
| dense_1 (Dense) | (None, 2) | 1026 |

=================================================================

Total params: 743,170

Trainable params: 743,170

Non-trainable params:

---

For 2D CNN on the graph representation, the architecture of network is as the following:

Layer (type) Output Shape Param #

```
=================================================================
conv2d_1 (Conv2D)            (None, 31, 31, 128)       1280
_________________________________________________________________
conv2d_2 (Conv2D)            (None, 29, 29, 128)       147584
_________________________________________________________________
max_pooling2d_1 (MaxPooling2 (None, 14, 14, 128)       0
_________________________________________________________________
conv2d_3 (Conv2D)            (None, 12, 12, 128)       147584
_________________________________________________________________
conv2d_4 (Conv2D)            (None, 10, 10, 128)       147584
_________________________________________________________________
max_pooling2d_2 (MaxPooling2 (None, 5, 5, 128)         0
_________________________________________________________________
conv2d_5 (Conv2D)            (None, 3, 3, 256)         295168
_________________________________________________________________
global_average_pooling2d_1 ( (None, 256)               0
_________________________________________________________________
dropout_1 (Dropout)          (None, 256)               0
_________________________________________________________________
dense_1 (Dense)              (None, 2)                 514
=================================================================
Total params: 739,714
Trainable params: 739,714
Non-trainable params: 0
_________________________________________________________________
```

Models were coded using Keras package (https://keras.io/), and trained using NVIDIA Tesla P4 GPU.

#### Database on CRISPR RNA processing

The parent vector contains an araC-GGGS-GFP fusion protein. The transcription is driven by the araC-repressed, arabinose-inducible promoter pBAD. The reporter plasmids were constructed by inserting a library of 33 nucleotides into GGGS linker, by Golden Gate assembly. The Golden Gate products were transformed into DH5a cells ($F^-$ *endA1 glnV44 thi-1 recA1 relA1 gyrA96 deoR nupG purB20* φ80d*lacZ*ΔM15 Δ(*lacZYA-argF*)U169, hsdR17($r_K^- m_K^+$), $\lambda^-$). The libraries of bacteria were induced by 0.2% arabinose, and then were sorted by GFP positive gate, to reduce the frequency of clones that are GFP negative due to trivial reasons such as stop codons. The enriched libraries were recovered under the same condition, and were again sorted for GFP positive strains in the same gate setting. After twice sorting, the distributions of GFP expression became unimodal. Then we made twice-sorted pools of GFP-positive cells competent, transformed Csy4-expressing plasmid. When the tested substrate is not cleaved by Csy4, araC-GFP is expressed; if the tested substrate is cleaved, GFP expression is repressed. In the presence of Csy4, the libraries are further sorted into 2 bins of GFP signal. Each group were then sorted again after overnight culture under the arabinose induction. We re-grew the bacteria from each groups, and measured their expression by flow cytometry, reproducing unimodal distributions. Overnight cultures of each group of library, were used to extract the plasmids. Inserted library is amplified, and sequenced by NextSeq 550 at Institut Cochin GENOM'IC platform. Then for each sequence detected, we calculated the ratio of reads in GFP negative and positive bins, and rounded the ratio into 0 or 1 labels.

#### Thermodynamic features

Full-length 33-nt sequences were analyzed by Nupack package (http://www.nupack.org/), to compute the partition function, complex energy of the whole RNA structural ensemble, and a 33 × 33 matrix of base-pairing probability in the thermodynamic equilibrium. Also using Nupack, the minimum free energy (mfe), mfe structure, and probability of RNA folding into mfe structure at the equilibrium were predicted. The total number of possible structures were also computed, and put together with other features.

#### RNA folding kinetic features

Using Kinwalker (https://www.tbi.univie.ac.at/RNA/kinwalker.1.html) in ViennaRNA package, we predicted the RNA folding trajectory. The intermediate structures, energy barrier and time required to get into the next structure, and the running time was recorded for each sequence. The statistics including mean, median, standard deviation, and maximum were applied on the energy barriers of each trajectory. Number of total intermediate structures were also counted.

#### Translational outcome

First, all the flanking sequences were translated into amino acid, and turned into integer encoding. Then the codon usages were listed according to the Genescript reference, which includes two representations, codon fraction in the same amino acid, and the codon frequency in 1,000 codons of all amino acids.

#### Base content, and abundances of k-mer, and ordered pairs

Bases content of each flanking context were computed, i.e. the percentage of ATGC in the flanking context. Remarkably, this feature is computed from each sequence. It’s different from the analysis on nucleotide content in each position which is conducted on the whole dataset. Abundancies of k-mer were listed for each sequence, for k from 2 to 4. For instance, the abundance of TT for sequence of GGATTTTGGAGGCCCCTT is 4. 16 2-mers, 4^3 = 64 3-mers, and 4^4 = 256 4-mers were considered. Matlab nmercount is used to count k-mers (https://ww2.mathworks.cn/help/bioinfo/ref/nmercount.html).

#### Mfe structure analysis

The mfe structures predicted before were used for detailed analyses. First, it'll be determined if the Csy4 binding site is folded into the 5-bp stem with 5-nt loop hairpin structure as desired. In the range from the upstream flanking context till the fifth base of binding site, the number of base that pairs to the downstream is computed, and vice versa. These two numbers should be the same, otherwise Csy4 binding site is not folded properly. If a stem loop larger including binding site exist, the total number of mismatches inside the stem loop were counted, which is the sum of bulge lengths. Then the adjacent flanking contexts were analyzed. If the adjacent flanking context forms in extra stems, a length of adjacent stem will be computed from binding site to the first mismatch. If the adjacent flanking context is single-stranded, a length of adjacent loop will be computed from binding site to the first base-pairing. All the computations mentioned above were computed separately for upstream and downstream flanking context, because their structural features are not always symmetric.

#### Single feature fitting

Individual features were used to fit 70% of the large dataset by logistic regression, using Matlab fitclinear function (https://ww2.mathworks.cn/help/stats/fitclinear.html). The resulting models were tested on 15% validation dataset, to estimate the predictive power of the features.

##### Random forest model

All the outcomes coming from the conventional analyses, including k-mer search, thermodynamic analysis, kinetic analysis, codon usage biases, and mfe structure analysis. Random forest model is created by Matlab TreeBagger (https://ww2.mathworks.cn/help/stats/treebagger.html) using 100 trees.

#### Data and code availability

Data used in this study, codes for neural networks, and models we acquired are available at https://github.com/maincover/sequence-dl-haotian
